# The two-speed genomes of filamentous pathogens: waltz with plants

**DOI:** 10.1101/021774

**Authors:** Suomeng Dong, Sylvain Raffaele, Sophien Kamoun

**Affiliations:** Department of Plant Pathology, Nanjing Agricultural University, Nanjing 210095, China; INRA, Laboratoire des Interactions Plantes-Microorganismes (LIPM), UMR441, Castanet-Tolosan 31326, France; CNRS, Laboratoire des Interactions Plantes-Microorganismes (LIPM), UMR2594, Castanet-Tolosan 31326, France; The Sainsbury Laboratory, Norwich Research Park, Norwich NR4 7UH, United Kingdom

**Author notes:** Corresponding author: Kamoun, Sophien.

## Abstract

Fungi and oomycetes include deep and diverse lineages of eukaryotic plant pathogens. The last 10 years have seen the sequencing of the genomes of a multitude of species of these so-called filamentous plant pathogens. Already, fundamental concepts have emerged. Filamentous plant pathogen genomes tend to harbor large repertoires of genes encoding virulence effectors that modulate host plant processes. Effector genes are not randomly distributed across the genomes but tend to be associated with compartments enriched in repetitive sequences and transposable elements. These findings have led to the “two-speed genome” model in which filamentous pathogen genomes have a bipartite architecture with gene sparse, repeat rich compartments serving as a cradle for adaptive evolution. Here, we review this concept and discuss how plant pathogens are great model systems to study evolutionary adaptations at multiple time scales. We will also introduce the next phase of research on this topic.

## Introduction

Infectious plant diseases wreak havoc to world agriculture and threaten renewed efforts to meet the food security needs of a booming world population. Among the most destructive plant pathogens are fungi and oomycetes, two groups of eukaryotes that are phylogenetically unrelated but exhibit similarities in their filamentous growth habit, heterotrophic lifestyle and specialized infection structures [1]. These so-called filamentous plant pathogens include notorious parasites, such as the rice blast fungus, several species of cereal-infecting rust fungi, the Irish potato famine pathogen, and many others that continue to trigger epidemics that threaten our food supply. Crop losses due to these filamentous pathogens amount to enough food to feed hundreds of millions of people [2]. Thus, it is not surprising that the plant pathology and genome biology communities teamed up to sequence the genomes of the main taxa of these formidable parasites. The first such genome, that of the rice blast fungus *Magnaporthe oryzae*, was reported only 10 years ago [3]. This was soon followed by the genomes of the oomycete soybean root rot pathogen *Phytophthora sojae* and the sudden oak death pathogen *P. ramorum* [4]. These landmark papers were accompanied by a flurry of publications with dozens of filamentous plant pathogen genomes becoming available for comparative analyses [5-12]. Interestingly, a number of concepts have emerged from comparative analyses of these genomes even though the species examined are highly diverse [13-19]. First, filamentous plant pathogen genomes tend to have distinctive architecture with an evolutionary trend towards large, repeat-bloated genomes. Second, these genomes harbor large repertoires of genes encoding secreted proteins, known as effector proteins, which function as virulence factors that modulate host plant processes. Finally, the effector genes are not randomly distributed across the genomes but tend to be associated with compartments enriched in repetitive sequences and transposable elements. These findings have led to the “two-speed genome” model in which pathogen genomes have a bipartite architecture with the gene sparse, repeat rich compartment serving as a cradle for adaptive evolution [5,17].

Remarkably, the unusual genome architecture and occurrence of effector genes in specific genome compartments is a feature that has evolved repeatedly in independent phylogenetic lineages of filamentous pathogens (**Figure 1**). This remarkable convergence in genome structure and its link to the pathogenic lifestyle of these organisms is a fundamental concept that has emerged from the last ∼10 years of comparative genome analyses of fungal and oomycete pathogens. Here, we review this concept and discuss how plant pathogens are great model systems to study evolutionary adaptations at multiple time scales. We will also introduce the next phase of research on this topic.

**Figure 1.**
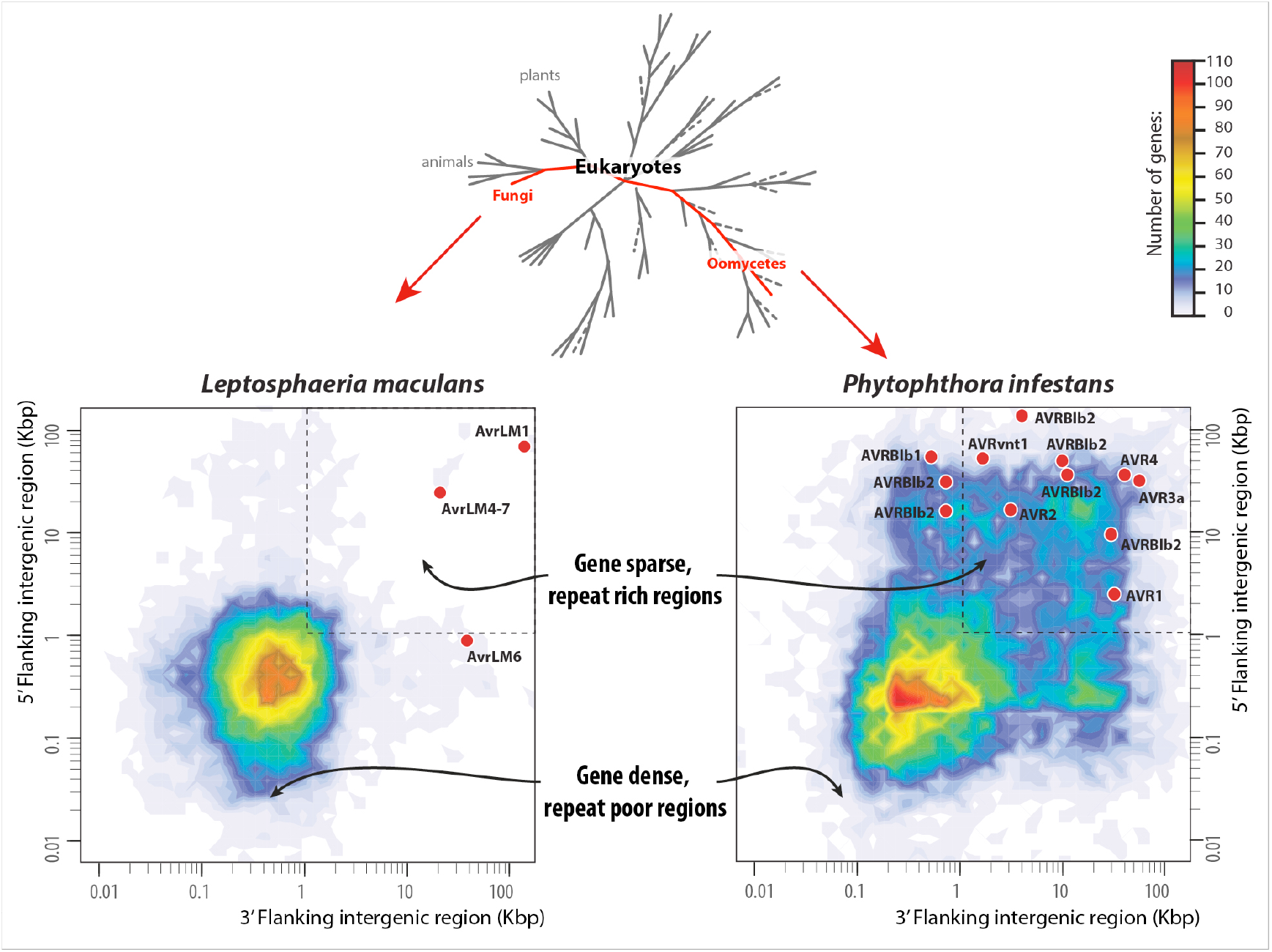
Convergence towards a similar genome architecture in phylogenetically unrelated plant pathogens. The flanking distance between neighbouring genes provides a measurement of local gene density and is displayed as a color-coded heat map based on a whole genome analysis of the fungus *Leptosphaeria maculans* and the oomycete *Phytophthora infestans* [23]. In addition, the figure displays the distribution of Avr effector genes of *L. maculans* and *P. infestans* according to the length of their 5’ and 3’ flanking intergenic regions. Note how in both cases the Avr effector genes primarily occupy the gene sparse regions of the genome.

### Genome size: bigger can be better

The textbook view is that parasites and symbionts tend to evolve smaller and more compact genomes than their free-living relatives [20,21]. Filamentous plant pathogens include some of the most glaring exceptions to this notion [5]. Although not all species have expanded genomes, several lineages of filamentous plant pathogens are characterized by a proliferation of repetitive DNA and markedly increased genome size. The trend applies to independent and ancient parasite lineages, including the ascomycete powdery mildew fungi (∼120 Mbp., 64% repetitive for *Blumeria graminis* DH14) and bamboo witches’ broom pathogen (∼60Mbp., 57% repetitive for *Aciculosporium take* MAFF241224), basidiomycete rust fungi (∼220 Mbp, 45% repetitive for *Melampsora lini* CH5), oomycetes of the genus *Phytophthora* (∼240 Mbp, 74% repetitive for *P. infestans* T30-4) and downy mildews (∼100 Mbp, 43% for *Hyaloperonospora arabidopsidis* Emoy2) [9,14,22-24]. In addition, several genomes, including those of the coffee rust and Asian soybean rust fungi, have not been fully sequenced yet because their large size and complexity have delayed these projects [25,26]. Convergence towards large genomes that are expanded compared to nonparasitic relatives is clearly an evolutionary trend in fungal and oomycete plant pathogens. But what are the evolutionary trade-offs of these repeat-bloated genomes?

### Two-speed genomes: strikingly uneven rates of evolution across filamentous pathogen genomes

A fundamental insight from analyses of filamentous plant pathogen genomes is that genes located in repeat-rich regions tend to evolve more rapidly than those in the rest of the genome [5,17]. This pattern could be demonstrated through comparative genomics of closely related species (**Figure 2**). Genome analyses of *P. infestans* and three of its sister species revealed uneven evolutionary rates across genomes with genes in repeat-rich regions showing higher rates of structural polymorphisms and positive selection [13]. A similar mosaic pattern was noted in the genome of the plant pathogenic fungus *Leptosphaeria maculans* [15,16]. Comparative genome analyses by Grandaubert et al. revealed that the expanded regions in the bipartite genome of Brassica infecting strains of *L. maculans* resulted from bursts of transposable element activity [16]. Genome analyses of the wheat fungal pathogen *Zymoseptoria tritici* and its closest known relatives also revealed a bipartite pattern of genome evolution with genes in essential and conditionally dispensable chromosomes evolving at strikingly different rates [27]. Comparison of complete genome sequences of 15 *Clavicipitaceae* fungi revealed frequent presence/absence polymorphisms and sequence variation in alkaloid biosynthesis gene clusters embedded in repeat-rich blocks [24,28]. These uneven patterns of genome evolution across filamentous pathogen genomes have led to the concept of “two-speed genome” with the gene sparse, repeat rich compartments serving as a cradle for adaptive evolution [5,17].

**Figure 2.**
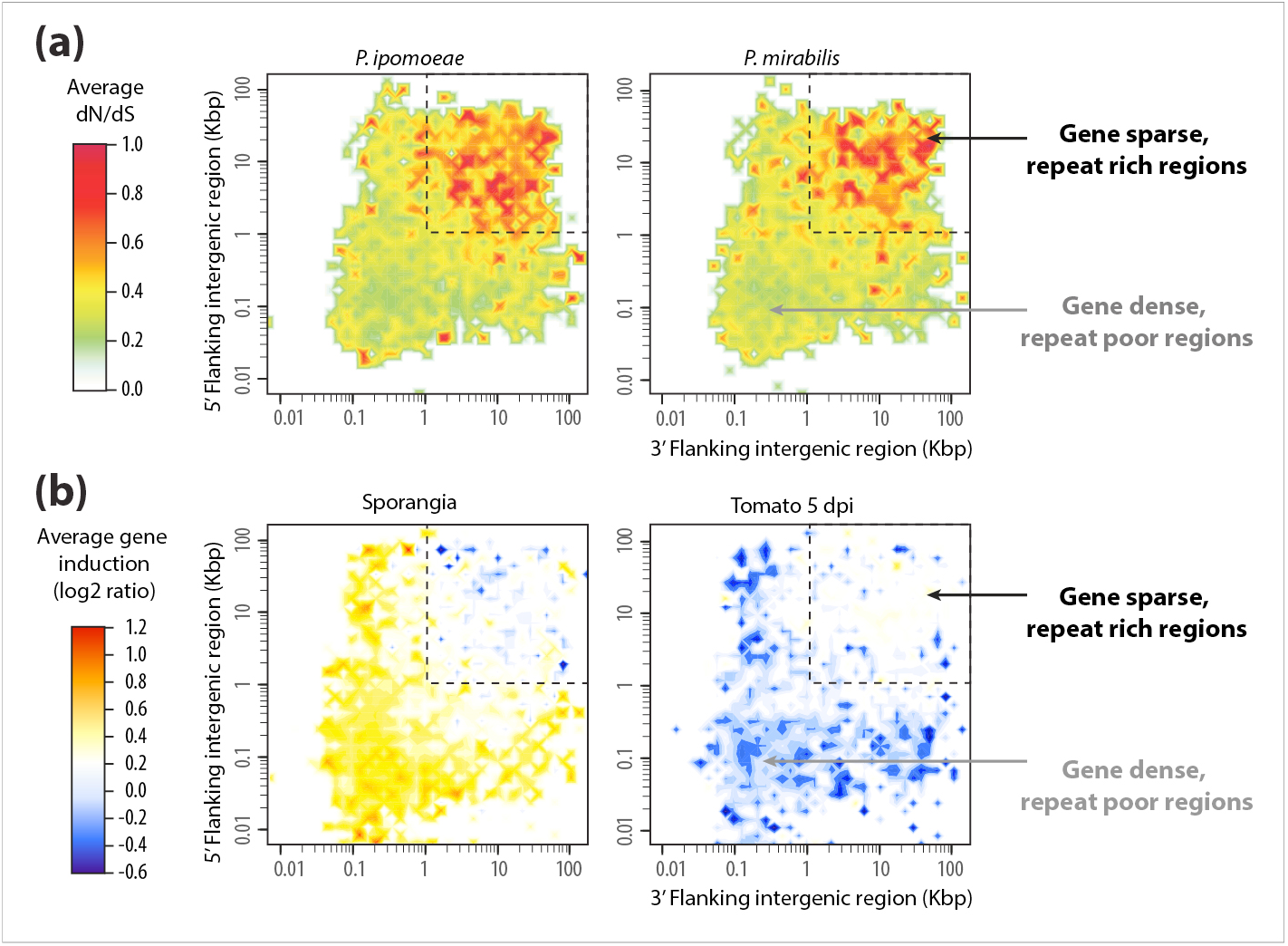
The two-speed genome of *Phytophthora infestans*. **(a)** Distribution of polymorphisms in *P. ipomoeae* and *P. mirabilis* according to local gene density (measured as length of 5’ and 3’ flanking intergenic regions) [13]. The average values of dN/dS ratio (nonsynonymous to synonymous substitution rate) associated with genes in each bin are shown as a color-coded heat map. The ratios were calculated relative to *P. infestans*, the sister species of *P. ipomoeae* and *P. mirabilis*. Note that the gene-sparse regions are highly enriched in genes under positive selection with elevated dN/dS ratio. **(b)** Distribution of gene expression induction in sporangia and during infection of tomato 5 days post inoculation according to local gene density. The average induction values associated to genes in each bin are shown as a color-coded heat map. Note that genes that are induced in sporangia, which are the asexual spores of *Phytophthora* spp., are enriched in the gene-dense compartments of the genome, however, the gene-sparse regions of *P. infestans* genome are highly enriched in plant-induced genes.

### Beyond comparative genomics: How does the two-speed genome accelerate pathogen evolution

Initially, repeat-rich, gene poor genome compartments of filamentous pathogens were primarily thought of as hot spots for duplication, deletion, and recombination that result in accelerated evolution through increased structural variation. However, it is becoming clear that there are more mechanisms by which this particular genome architecture enables a faster tempo of pathogen evolution [5,17,18,29]. In some ascomycete fungi, close proximity of effector genes to degenerated transposable elements can result in increased levels of local mutagenesis through repeat-induced point (RIP) mutations. In both France and Australia, effector alleles of *L. maculans* with premature stop codons or non-synonymous RIP mutations have evolved to evade plant resistance mediated by the RLM immune receptors in just a few years [30-32]. Epigenetic silencing could also leak from TEs to neighboring genes thus disproportionally affecting genes in the repeat-rich regions of the genome [33-35]. Proximity to transposable elements may have enabled the persistence of the *M. oryzae* effector *Avr-Pita* in asexual populations of the rice blast fungus *M. oryzae* despite repeated adaptive deletions that generated genotypes virulent on rice plants carrying the *Pi-ta* resistance gene [36]. The occurrence of this “mobile effector” in different chromosomes of *M. oryzae* and related *Pyricularia* species is consistent with its lateral transfer through parasexual genetic exchange, a process that probably involves hitchhiking with a neighboring retrotransposon. Horizontal gene transfer can also be facilitated by accessory chromosomes, which are known to move laterally in fungi following hyphal anastomosis [19].

### Jump or die: convergence to the two-speed genome architecture might be driven by macroevolutionary dynamics

Several oomycetes and fungi belong to deep lineages of plant pathogens. That they have engaged in uninterrupted antagonistic coevolutionary conflict with plants and yet managed to persist in the biota over millions of years is truly astonishing. The key to this evolutionary success is the ability of these fickle pathogens to overcome plant immunity, adapt to resistant plant varieties, and occasionally jump to new host species [5]. The distinct patterns of two-speed genome architecture identified in unrelated lineages of filamentous plant pathogens is intimately linked to the rapid evolutionary tempo of their effector genes and is, therefore, directly impacting their pathogenic lifestyle (**Figure 2B**). This remarkable convergence in genome architecture is best explained through the concept of clade selection. Lineages that evolved two-speed genomes have increased survivability – they are less likely to go extinct compared to lineages with less adaptable genomes, which are more likely to be purged out of the biota as their hosts develop full resistance or become extinct. In this “jump or die” model, pathogen lineages that have an increased likelihood to produce virulent genotypes on resistant hosts and non-hosts benefit from a macroevolutionary advantage and end up dominating the biota [5].

### Hidden in the genomes: diverse and large effector repertoires

Genome sequencing of filamentous plant pathogen genomes has typically revealed large repertoires of (candidate) effector genes. This trend is particularly marked in oomycete pathogens. In *Phytophthora*, downy mildews, and *Albugo* spp. effector genes can be readily annotated in genome sequences because they contain signal peptides followed by conserved motifs, e.g. RXLR in *Phytophthora* and downy mildews [37]. The number of predicted effector genes is staggering. In the potato blight pathogen *P. infestans*, ∼550 RXLR effector genes have been detected [23]. Another effector family, the Crinklers or CRNs, has also dramatically expanded in *P. infestans* to ∼200 genes and ∼250 pseudogenes [23]. Similar expanded secretomes have also been described in fungal plant pathogens. About 400-500 candidate secreted effector proteins (CSEPs) have been identified in the genomes of the barley and wheat powdery mildew fungus *B. graminis* [6,7,14]. In the rust fungi, up to 725 candidate effectors have been reported [9,38-40]. These effectors are thought to have a diversity of activities inside plant cells with several studies revealing that they target multiple host compartments and processes [41-44]. Effectors can also be metabolites that are coded by biosynthetic enzymes [45]. In *Clavicipitaceae* fungi, alkaloids generally serve as metabolite effectors produced by these plant-associated fungi to protect their host plants from predators [28]. Alkaloid biosynthesis loci are unstable due to surrounding large and pervasive blocks of repeats, thus increasing chemotypic variation of the alkaloids [24].

### The impact of host jumps on pathogen evolution

Several filamentous plant pathogens have evolved by shifting or jumping from one host plant to another [46-48]. In some cases, the jumps are to plants that are distantly related to the ancestral host. Such events are expected to dramatically impact effector evolution considering that these proteins function inside a plant cellular environment and target plant molecules and processes (**Figure 3**). In case of host jumps to distant plant species, a subset of the effector repertoire may become obsolete because the effector target may not be absent in the new host. Indeed, effector gene deletion is rampant in the genome of the smut fungus *M. pennsylvanicum*, which is thought to have evolved from a grass pathogen into a dicot pathogen [49]. Similarly, an elevated rate of copy number variation was detected among effector genes of *P. infestans* and its sister species, which evolved by jumps to hosts from unrelated botanical families [13]. In other cases, effectors may adapt to a different target in the new host resulting in the accumulation of mutations that improve or expand the activity of the effector. Consistent with this model, a large number of effector and other *in planta* induced genes displays signatures of adaptive evolution among the *Phytophthora* clade 1c species [13,50].

**Figure 3.**
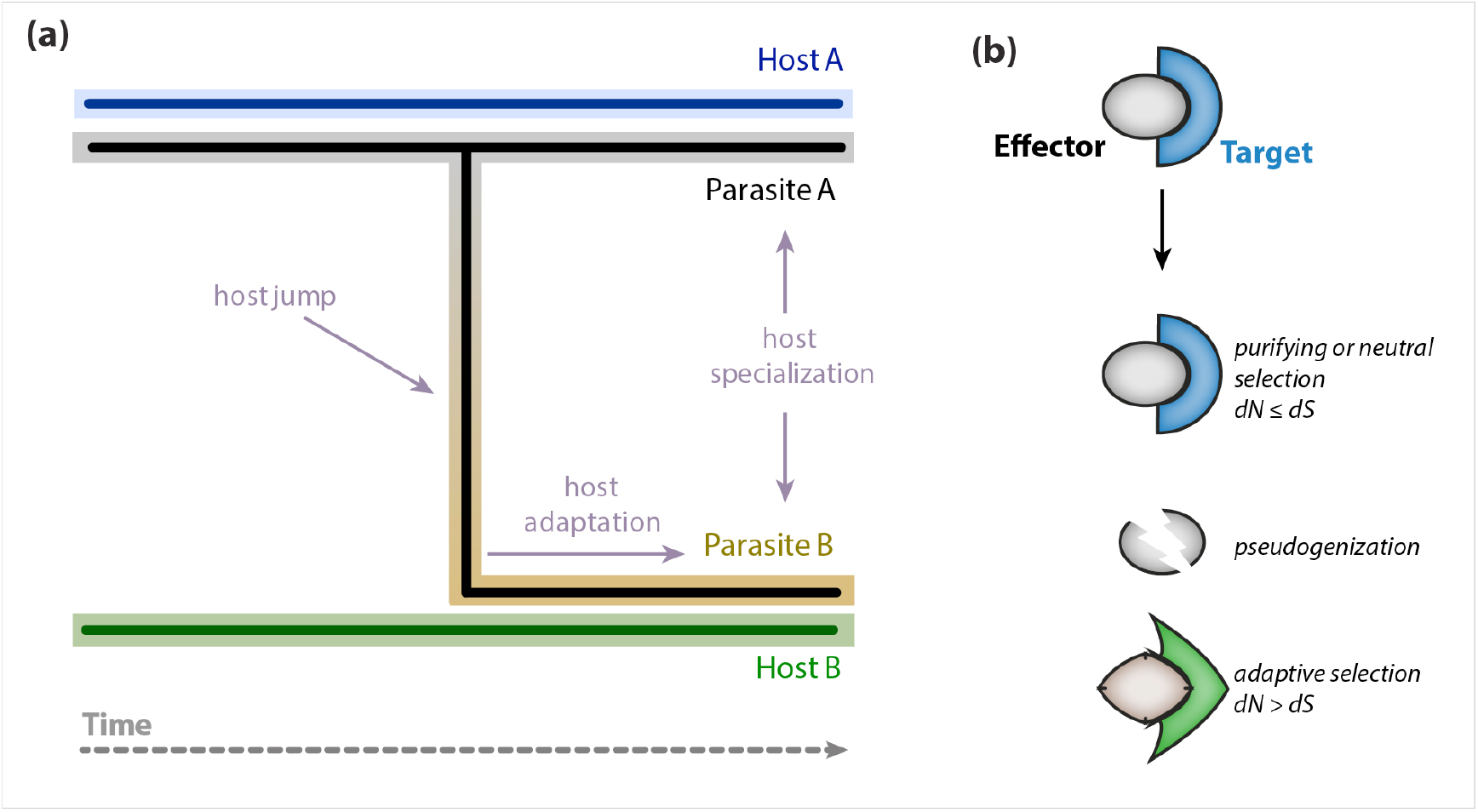
Impact of host jumps on effector evolution. **(a)** Schematic view of a hypothetical host jump of parasite A to host B followed by adaptation, specialization and eventually speciation of parasite B. The host jump is expected to have a dramatic effect on the effector repertoire of parasite A as effectors become exposed to a different host environment. The scheme in **(b)** depicts the three major scenarios of effector evolution following host jumps. We hypothesize that effector evolution in parasite B will be driven by the nature of the effector targets in the new host B. Panel **(a)** was adapted from Roy [46].

### Host adaptation and specialization as determinants of effector evolution

Filamentous fungi and oomycetes provide striking examples of rapid evolutionary adaptations. These are particularly well documented for effectors that have a so-called “avirulence” (Avr) activity, namely that trigger receptor-mediated plant immunity [51,52]. Avr effector genes have evolved a variety of allelic variants to evade plant immunity. In some cases, these variants carry amino acid polymorphisms that evade activating host immunity while presumably preserving the effector’s virulence activity. An example is the AVR-Pik gene of the rice blast fungus *M. oryzae*, which occurs as an allelic series with only a few amino acid polymorphisms that are mirrored by the matching resistance protein Pik-1 [53]. These polymorphisms appear to reflect arms race coevolutionary dynamics between *M. oryzae* and rice, with resistance responses correlating with binding of AVR-Pik to Pik-1 [53]. In other cases, gain of virulence was achieved through loss of function mutations such as premature stop codons or deletions [32,54]. Daverdin et al. monitored over four years the evolution of *L. maculans* populations faced with newly released oilseed rape cultivars with the *Rlm7* resistance gene [32]. *L. maculans* isolates virulent on *Rlm7* plants carried a diversity of independent molecular events in the matching effector gene AvrLm4-7, with RIP mutations and deletions the most frequent events [32]. Transcriptional silencing is yet another mechanism by which Avr effectors can evade host resistance [35,55]. Silencing of *P. sojae* effector gene *Avr3a* enables this pathogen to gain virulence on soybean plants carrying the resistance gene *Rps3* [33]. Silenced *Avr3a* alleles were transmitted to progeny in non-Mendelian fashion and persisted over multiple generations suggesting that epigenetic changes at this locus mediated the gain of virulence phenotype [33].

Effectors can also adapt to their host target in order to better manipulate host defense. In one case, the biochemical basis of effector adaptation following a host jump could be resolved [50]. *P. infestans* cystatin-like effector EPIC1 and its ortholog from the sister species *P. mirabilis* PmEPIC1 are more effective inhibitors of proteases from their respective hosts, RCR3 from potato and MRP2 from *Mirabilis jalapa*, respectively [50]. A Gln to Arg substitution at position 111 in PmEPIC1, a derived mutation unique to the *P. mirabilis* lineage, increased MRP2 inhibition. However, this mutation carries a trade-off as it impairs the effector’s ability to inhibit RCR3 proteases. This provides a molecular framework for effector specialization. An effector that evolved higher activity on a target in its new host performs poorly on the presumed ancestral host target. Evolution can therefore drive functional specialization of effectors during plant-pathogen coevolution, ultimately leading to the emergence and diversification of pathogen species [50].

## Outlook

Pathogenomics of fungi and oomycetes is an emerging field of research in plant pathology. Many of these pathogens belong to deep lineages that have successfully engaged in antagonistic coevolution with plants over millions of years. Genome sequencing of unrelated taxa revealed a remarkable convergence in several independent lineages towards similar genome architectures with effector genes populating repeat-rich genome compartments. “Two-speed” genome architectures may have emerged as a successful solution to survivability of these pathogen lineages in the face of an ever-changing host environment [56]. Although more pathogen genomes need to be sequenced and analyzed, research is edging towards a mechanistic understanding of filamentous pathogen genome evolution. How two-speed genomes accelerate pathogen evolution and counteract lineage extinction is poised to be a key question for the next 10 years.

Plant pathogens are great model system to study rapid evolutionary adaptations. In many cases, particularly with regards to Avr effector-immune receptor dynamics, the genetic determinants and the nature of the adaptive mutations are known [see for example 57]. However, effector adaptation to host targets is not as well understood as coevolution with immune receptors. We need to gain a better view of the tempo of effector coevolution with host factors other than immune receptors.

We still know little about how pathogens and their effectors adapt and specialize on new hosts. We also need to understand better the determinants of host specificity. How do broad host range pathogens cope with their multiple hosts? How are their genomes shaped by their interactions with multiple hosts? An example of a genomics-driven insight is a recent study on the generalist parasite *Albugo candida*, which consists of multiple races that specialize on particular Brassicaceae species [58]. The genomes of host-specialized *A. candida* races display a mosaic architecture with evidence of genetic exchanges. Host immunosuppression by one *A. candida* race enables colonization by another race therefore explaining how genetic exchange could take place between specialized genotypes with non-overlapping host ranges. This led McMullan et al. to propose that the immunosuppression capacity of *A. candida* enables this species to maintain a wide host-range through occasional inter-race genetic exchanges and reassortment of effector gene repertoires [58]. Ultimately, this process enables *A. candida* populations to counteract the trade-offs associated with host-specialization and persist over long evolutionary times.

In conclusion, we need to capitalize on the conceptual advances in the field of plant-microbe interactions to expand our understanding of host-pathogen evolution. Future studies ought to combine comparative genomics analyses with mechanistic and biochemical studies of virulence in order to gain a thorough understanding of filamentous pathogen adaptation to their hosts across multiple evolutionary time scales.

## Acknowledgements

We are thankful to past and present members of the Kamoun Lab as well as to several colleagues for numerous discussions and ideas. This review was developed based on reference [5] and was adapted from a talk available on YouTube (http://youtu.be/kogoAS_9Bgk). SD receives funding support from Chinese National Science Fund for Excellent Young Scholar (31422044). SR is supported by the European Research Council (ERC-StG 336808) and the Laboratory of Excellence project TULIP (ANR-10-LABX-41; ANR-11-IDEX-0002-02). SK receives funding primarily from the Gatsby Charitable Foundation, Biotechnology and Biological Sciences Research Council (BBSRC, UK), and European Research Council (ERC-NGRB).

*of special interest

**of outstanding interest

